# Reconstructing the Phylogeny of *Calliandra* sect. *Androcallis* (Fabaceae): Insights from the Inclusion of Colombian Species, with a Focus on the Enigmatic Taxon *Calliandra medellinensis*

**DOI:** 10.1101/2023.10.02.560511

**Authors:** Tatiana Arias, Juan David Saldarriaga, Henry Arenas-Castro, Álvaro Idárraga-Piedrahita, Norberto López-Alvarez, Eduardo Tovar Luque, Germán Torres-Morales, Mailyn A. Gonzalez, Iván Darío Soto-Calderón

## Abstract

Phylogenetic relationships for the genus *Calliandra* section *Androcallis* (Fabaceae) were reconstructed, including previously sequenced species from Central and South America and unexamined species from Colombia, one of *Calliandra* main diversity centers. Here, we generated novel DNA sequences of *Calliandra* species from Colombia for the nuclear Internal Transcribed Spacer (*ITS)* and the chloroplast *trnL* and *trnL-F* intergenic spacer. By incorporating a broader taxonomic sampling, the relationships among main clades in *Androcallis* were clarified, providing a systematics framework in which to test evolutionary hypotheses. Phylogenetic analysis recovered five well-supported clades within *Androcallis*. Most species within each clade had similar geographical distributions and relationships between the five major clades are strongly supported for the first time. However, core *Androcallis* relationships, including most species from Colombia sequenced here, remain unclear. A second goal of this study was to determine the taxonomic status of *Calliandra medellinensis*. This enigmatic taxon emblematic of Medellín, Colombia, is found in limited numbers within the Aburrá Valley and has been proposed to be a hybrid taxon. Here, *C. medellinensis*, *C. magdalenae* and *C. haematocephala* were not monophyletic within the core *Androcallis* clade. This suggests that *C. medellinensis* could potentially be an interspecific hybrid between *C. magdalenae* and *C. haematocephala*, thus challenging the taxonomic status of this species; however, more informative molecular markers should be used in future studies. Specifically, genomic studies should assess interspecific hybridization demographic models. Such insights can illuminate the *C. medellinensis* origin, guiding conservation strategies and providing valuable evolutionary overviews.

## INTRODUCTION

The *Calliandra* genus (subfamily Caesalpinioideae, in the Mimosoid clade of Fabaceae) comprises around 130 species of mostly shrubs and small trees naturally thriving in seasonally dry tropical forests, savannahs, and open rock fields (Barneby 1998; Bello and Forero 2005; LPWG 2017). Species occur in three main centers of diversity: northern Colombia and Venezuela, the Mexican Plateau, and the Northeast Region of Brazil (Barneby 1998; Souza and Queiroz 2004; Lewis et al. 2005; Souza et al. 2013). Plagiotropic branches of *Calliandra* trees offer shade from the sun and for that reason are commonly planted in urban spaces such as parks and gardens across South American towns and cities. Beyond their horticultural significance, these species have utility in construction, forage, and charcoal production, from which the Spanish term “carbonero” originates. Moreover, they play a crucial ecological role by fixing nitrogen into their roots in rhizobia/nodules (Bello and Forero 2005; Simpson 2006).

Bentham (1840) initially established the genus based on floral characteristics like monadelphous androecium with multiple stamens and fruits exhibiting elastic dehiscence from the apex. This classification evolved in 1844, with Bentham recognizing 60 species across five series, primarily distinguished by leaf and inflorescence morphology. Subsequent revisions incorporated Old-World species into the genus (Bentham 1875; Harms 1921; Paul 1979; Thulin et al. 1981). However, Barneby’s work (1998) restricted *Calliandra* to the New World for taxonomic reclassification. He introduced palynological traits uniting all species, including calymmate polyads with eight pollen grains connected by a common exine and viscous appendage on the basal cell that facilitates adherence to stigma surfaces during pollination (Guinet 1965; Prenner and Teppner 2005; Greissl 2006). Barneby (1998) further proposed a revised taxonomy, introducing five sections and 14 series within the genus based on inflorescence structure: *Calliandra* sect. *Calliandra* featuring terminal pseudo-racemes of heads or umbels; *C.* sect. *Microcallis* with heads or umbels arising from short lateral branches; sect. *Androcallis* with a terminal umbel, and *C.* sect. *Acistegia* with lateral inflorescences bearing stipular spicules and sect. *Acroscias* a monotypic species with a highly specialized single terminal umbel.

Despite these advances, Barneby (1998) excluded the Old-World taxa from *Calliandra sensu stricto*. The most recent phylogeny by Souza et al. (2013) relied on the Bentham (1840) and Barneby (1998) classifications, identifying *Calliandra* as monophyletic based on nuclear *ITS* and chloroplast *trnL-F* regions. They introduced a novel infrageneric classification, unveiling three new sections: *Calliandra* sect. *Tsugoideae* (four species from northwestern South America), *C.* sect. *Septentrionales* (six species in arid areas of the United States and Mexico), and *C.* sect. *Montila* (37 species restricted to Brazil). However, Souza et al. (2013) lacked samples from Colombian, a pivotal diversity center for the genus, hindering a comprehensive understanding of *Calliandra* evolutionary relationships.

The *Calliandra* species occurring in Colombia comprise 22 taxa, including widespread species like *C. haematocephala* and *C. magdalenae*, as well as regionally confined species such as *C. antioquiae* and *C. medellinensis* (Fig. 1) (Idárraga et al. 2011; Bernal et al. 2016). In particular, the endemic *Calliandra medellinensis*, thriving within the metropolitan expanse of Medellín (The Aburrá Valley), merits attention because of its shrubby habit, glabrous branches, and inflorescences bearing solitary heads with seven to eight flowers reds (Bello and Forero 2005). Although described by Britton and Killip in 1936, its true taxonomic status remains ambiguous, with suggestions of intermediate traits between *C. haematocephala* and *C. magdalenae* potentially involving hybrid origins (Barneby 1998, Bello and Forero 2005), aspects yet unexplored using molecular markers.

**Figure 1.**
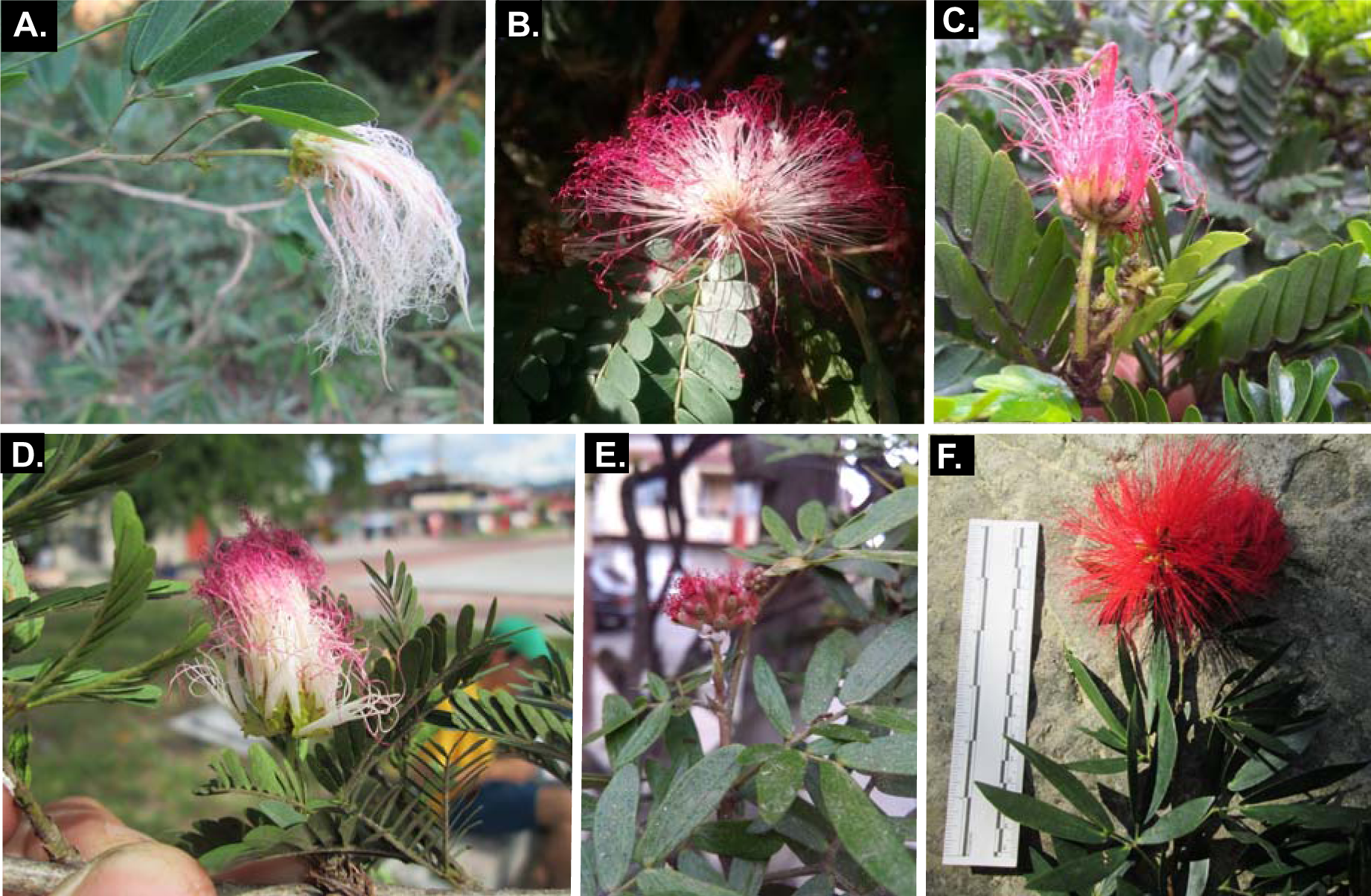
Representative vegetative and floral diversity of some *Calliandra* species that occur in Colombia. A. *Calliandra tergemina*. B. *Calliandra magdalenae*. C. *Calliandra medellinensis*. D. *Calliandra riparia*. E. *Calliandra falcata*. F. *Calliandra antioquiae*.

This study incorporates 12 *Calliandra* species present in Colombia into the existing dataset of Souza et al. (2013) leveraging nuclear *ITS* and chloroplast *trnL-F* sequences. By including Colombian species, we aimed to elucidate *Calliandra* systematics and resolve *C. medellinensis* systematic position and species status, while assessing its conservation urgency following IUCN guidelines (2018).

## METHODS

### Plant material and collections

To ascertain the *Calliandra* species present in Colombia and their distributions, we conducted an extensive review of bibliographic records and examined botanical specimens from various herbaria, including the Joaquín Antonio Uribe at Medellín Herbarium (JAUM), National University Herbarium at Medellín (MEDEL), University of Antioquia Herbarium (HUA), Colombian National Herbarium (COL), and data sourced from the Global Biodiversity Information Facility (GBIF), Urban Tree System (SAU) of Medellín, and Catalogue of Plants and Lichens of Colombia (Bernal et al. 2016).

To acquire tissue samples from the identified species, we accessed *Calliandra* samples archived at the Humboldt Institute’s Biological Tissues Collection (IAVH-CT) and conducted fieldwork across three departments in Colombia (Antioquia, Tolima, and Cundinamarca) (Fig. 2), collecting leaf tissue from the remaining species (González and Arenas-Castro 2017). The collected tissue samples were deposited at IAVH-CT, whereas the corresponding voucher specimens were lodged at JAUM, HUA, and the Humboldt Institute Herbarium (FMB) (Table S1).

**Figure 2.**
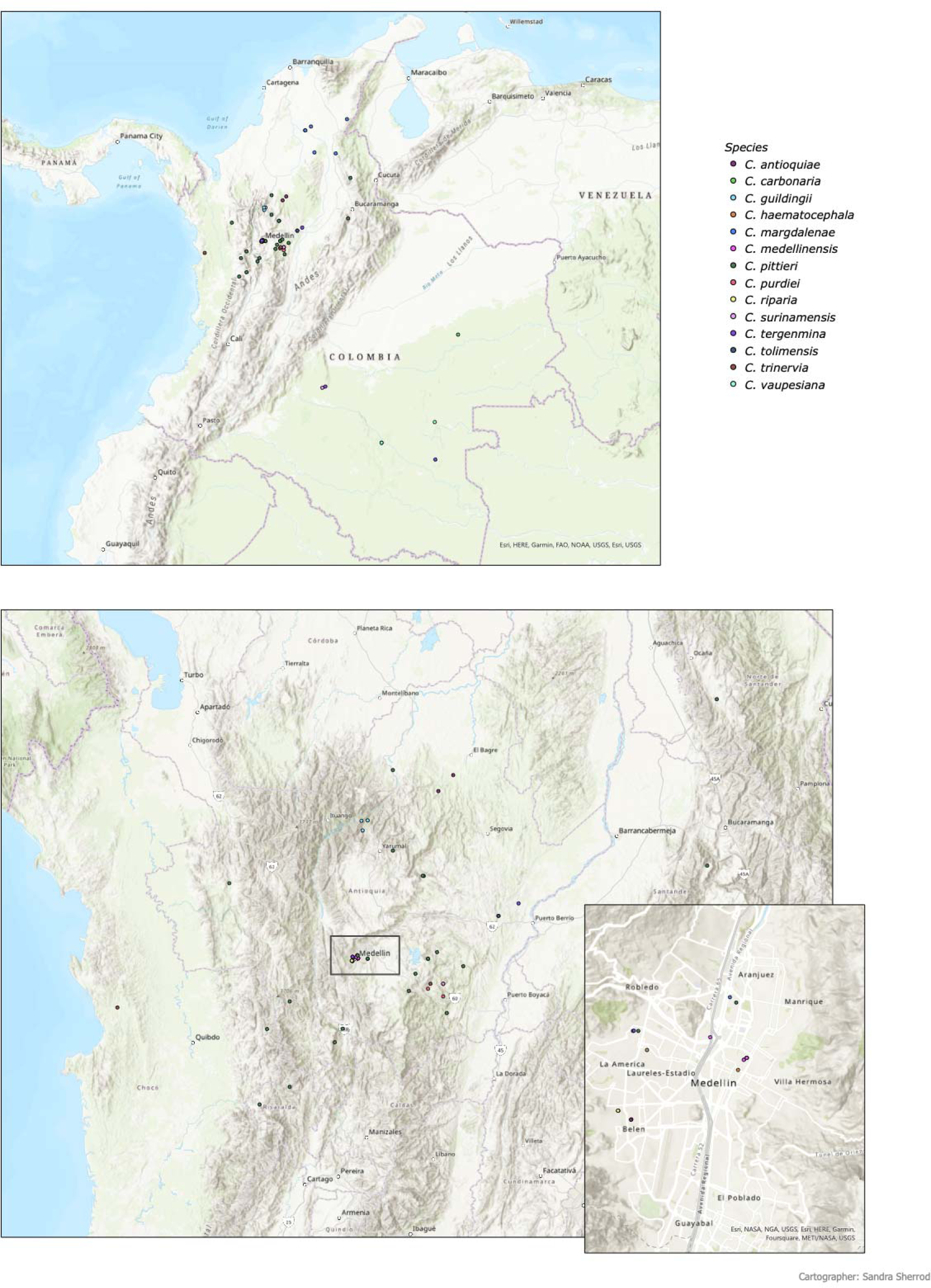
Distribution of *Calliandra* species in Colombia based on herbarium specimens.

Our sampling encompassed 27 documented *C. medellinensis* specimens, along with specimens from 12 of the 22 *Calliandra* species previously reported in Colombia (Fig. 1, Table 1-2). Species such as *Calliandra antioquiae* Barneby, *C. tolimensis* Taub., and *C. medellinensis* Britton and Killip were novel inclusions within *Calliandra* phylogeny. The remaining species were part of earlier phylogenetic studies, although representatives of Colombia were absent. Species identification adhered to a taxonomic key tailored for *Calliandra* species in Colombia (Forero and Romero 2005).

**Table 1.**
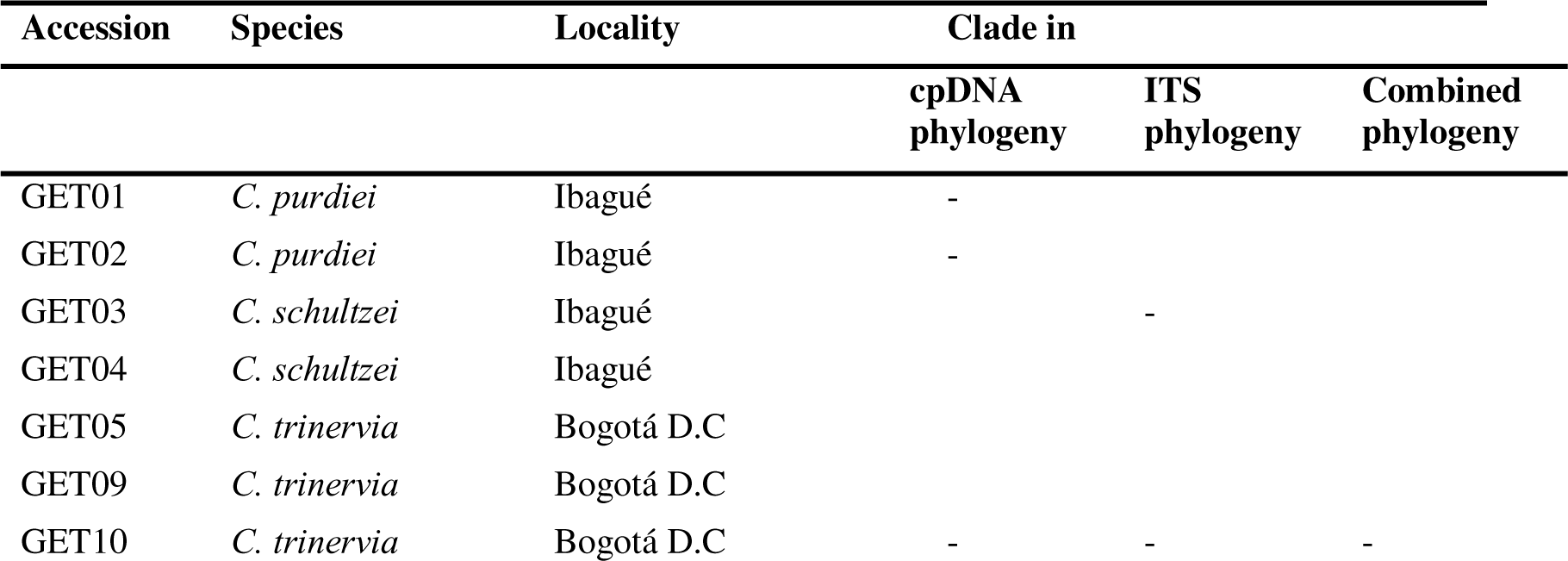
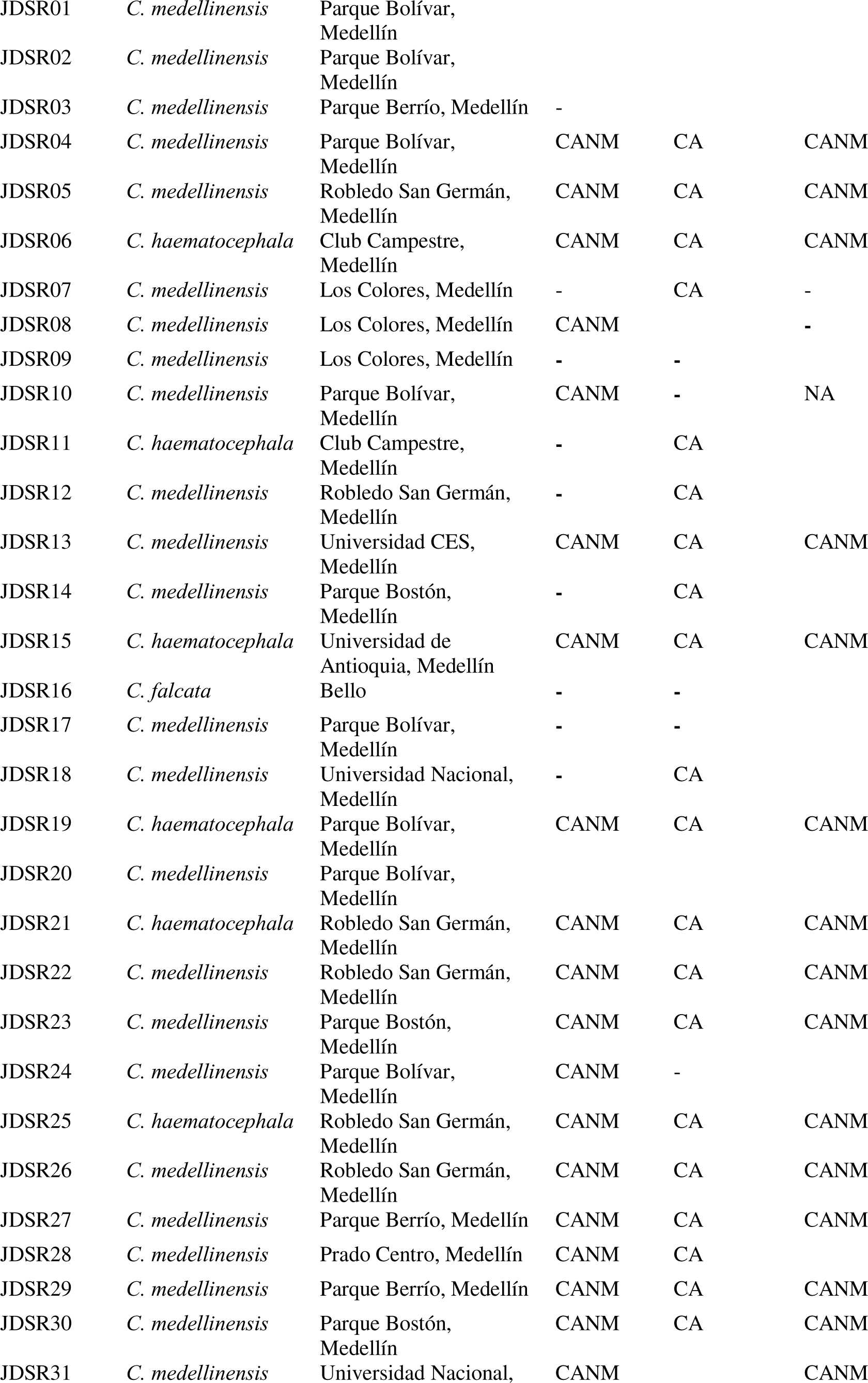
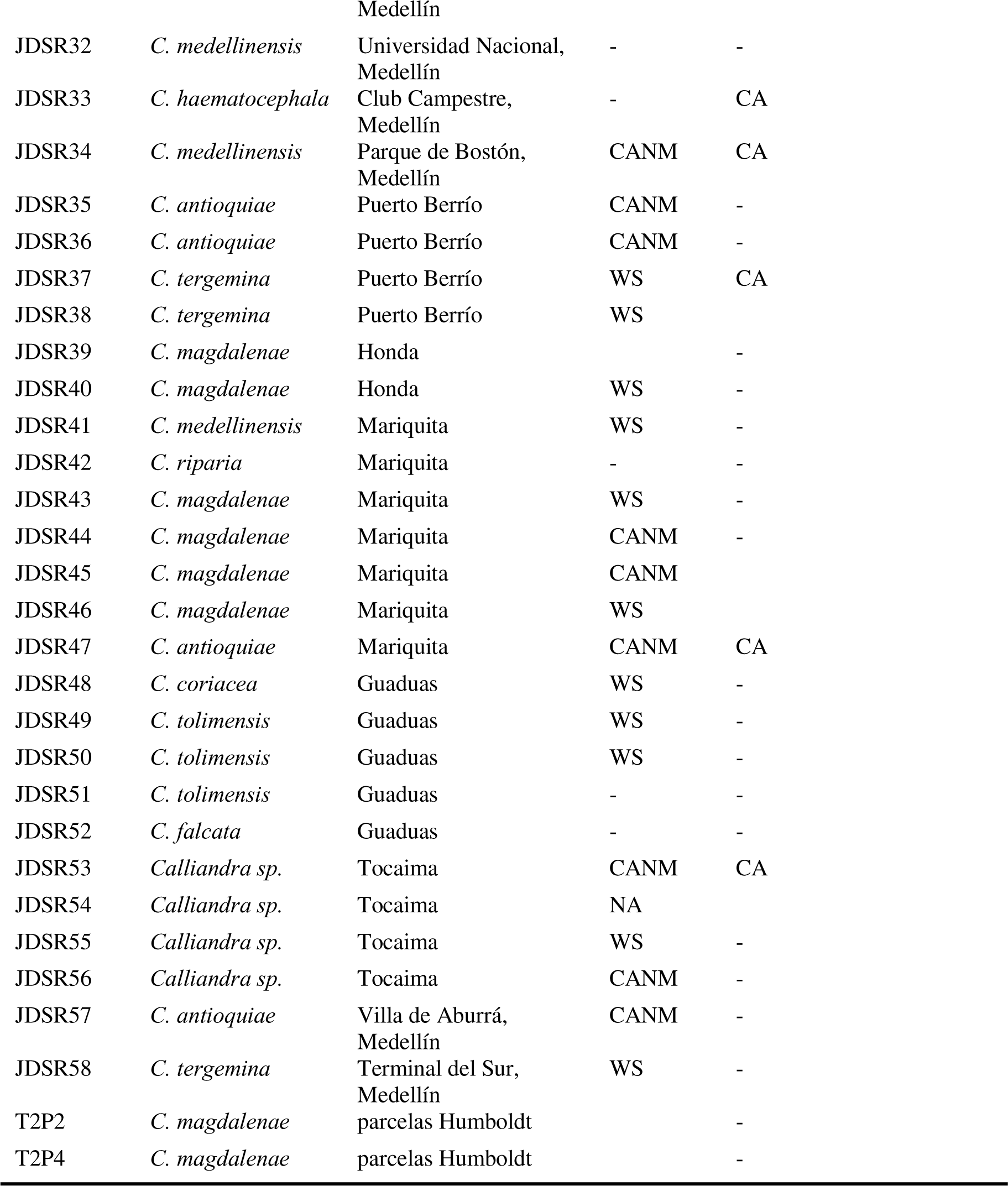
Accessions of *Calliandra* collected in Colombia and their phylogenetic position. CA: Core Androcallis; CANM: Core Androcallis non monophyletic.

| Accession | Species | Locality | Clade in |  |  |
| --- | --- | --- | --- | --- | --- |
|  |  |  | cpDNA<br>phylogeny | ITS<br>phylogeny | Combined<br>phylogeny |
| GET01 | *C. purdiei* | Ibagué | - |  |  |
| GET02 | *C. purdiei* | Ibagué | - |  |  |
| GET03 | *C. schultzei* | Ibagué |  | - |  |
| GET04 | *C. schultzei* | Ibagué |  |  |  |
| GET05 | *C. trinervia* | Bogotá D.C |  |  |  |
| GET09 | *C. trinervia* | Bogotá D.C |  |  |  |
| GET10 | *C. trinervia* | Bogotá D.C | - | - | - |
| JDSR01 | *C. medellinensis* | Parque Bolívar,<br>Medellín |  |  |  |
| JDSR02 | *C. medellinensis* | Parque Bolívar,<br>Medellín |  |  |  |
| JDSR03 | *C. medellinensis* | Parque Berrío, Medellín | - |  |  |
| JDSR04 | *C. medellinensis* | Parque Bolívar,<br>Medellín | CANM | CA | CANM |
| JDSR05 | *C. medellinensis* | Robledo San Germán,<br>Medellín | CANM | CA | CANM |
| JDSR06 | *C. haematocephala* | Club Campestre,<br>Medellín | CANM | CA | CANM |
| JDSR07 | *C. medellinensis* | Los Colores, Medellín | - | CA | - |
| JDSR08 | *C. medellinensis* | Los Colores, Medellín | CANM |  | - |
| JDSR09 | *C. medellinensis* | Los Colores, Medellín | - | - |  |
| JDSR10 | *C. medellinensis* | Parque Bolívar,<br>Medellín | CANM | - | NA |
| JDSR11 | *C. haematocephala* | Club Campestre,<br>Medellín | - | CA |  |
| JDSR12 | *C. medellinensis* | Robledo San Germán,<br>Medellín | - | CA |  |
| JDSR13 | *C. medellinensis* | Universidad CES,<br>Medellín | CANM | CA | CANM |
| JDSR14 | *C. medellinensis* | Parque Bostón,<br>Medellín | - | CA |  |
| JDSR15 | *C. haematocephala* | Universidad de<br>Antioquia, Medellín | CANM | CA | CANM |
| JDSR16 | *C. falcata* | Bello | - | - |  |
| JDSR17 | *C. medellinensis* | Parque Bolívar,<br>Medellín | - | - |  |
| JDSR18 | *C. medellinensis* | Universidad Nacional,<br>Medellín | - | CA |  |
| JDSR19 | *C. haematocephala* | Parque Bolívar,<br>Medellín | CANM | CA | CANM |
| JDSR20 | *C. medellinensis* | Parque Bolívar,<br>Medellín |  |  |  |
| JDSR21 | *C. haematocephala* | Robledo San Germán,<br>Medellín | CANM | CA | CANM |
| JDSR22 | *C. medellinensis* | Robledo San Germán,<br>Medellín | CANM | CA | CANM |
| JDSR23 | *C. medellinensis* | Parque Bostón,<br>Medellín | CANM | CA | CANM |
| JDSR24 | *C. medellinensis* | Parque Bolívar,<br>Medellín | CANM | - |  |
| JDSR25 | *C. haematocephala* | Robledo San Germán,<br>Medellín | CANM | CA | CANM |
| JDSR26 | *C. medellinensis* | Robledo San Germán,<br>Medellín | CANM | CA | CANM |
| JDSR27 | *C. medellinensis* | Parque Berrío, Medellín | CANM | CA | CANM |
| JDSR28 | *C. medellinensis* | Prado Centro, Medellín | CANM | CA |  |
| JDSR29 | *C. medellinensis* | Parque Berrío, Medellín | CANM | CA | CANM |
| JDSR30 | *C. medellinensis* | Parque Bostón,<br>Medellín | CANM | CA | CANM |
| JDSR31 | *C. medellinensis* | Universidad Nacional, | CANM |  | CANM |

|  |  |  |  |  |
| --- | --- | --- | --- | --- |
|  |  | Medellín |  |  |
| JDSR32 | *C. medellinensis* | Universidad Nacional,<br>Medellín | - | - |
| JDSR33 | *C. haematocephala* | Club Campestre,<br>Medellín | - | CA |
| JDSR34 | *C. medellinensis* | Parque de Bostón,<br>Medellín | CANM | CA |
| JDSR35 | *C. antioquiae* | Puerto Berrío | CANM | - |
| JDSR36 | *C. antioquiae* | Puerto Berrío | CANM | - |
| JDSR37 | *C. tergemina* | Puerto Berrío | WS | CA |
| JDSR38 | *C. tergemina* | Puerto Berrío | WS |  |
| JDSR39 | *C. magdalenae* | Honda |  | - |
| JDSR40 | *C. magdalenae* | Honda | WS | - |
| JDSR41 | *C. medellinensis* | Mariquita | WS | - |
| JDSR42 | *C. riparia* | Mariquita | - | - |
| JDSR43 | *C. magdalenae* | Mariquita | WS | - |
| JDSR44 | *C. magdalenae* | Mariquita | CANM | - |
| JDSR45 | *C. magdalenae* | Mariquita | CANM |  |
| JDSR46 | *C. magdalenae* | Mariquita | WS |  |
| JDSR47 | *C. antioquiae* | Mariquita | CANM | CA |
| JDSR48 | *C. coriacea* | Guaduas | WS | - |
| JDSR49 | *C. tolimensis* | Guaduas | WS | - |
| JDSR50 | *C. tolimensis* | Guaduas | WS | - |
| JDSR51 | *C. tolimensis* | Guaduas | - | - |
| JDSR52 | *C. falcata* | Guaduas | - | - |
| JDSR53 | *Calliandra sp.* | Tocaima | CANM | CA |
| JDSR54 | *Calliandra sp.* | Tocaima | NA |  |
| JDSR55 | *Calliandra sp.* | Tocaima | WS | - |
| JDSR56 | *Calliandra sp.* | Tocaima | CANM | - |
| JDSR57 | *C. antioquiae* | Villa de Aburrá,<br>Medellín | CANM | - |
| JDSR58 | *C. tergemina* | Terminal del Sur,<br>Medellín | WS | - |
| T2P2 | *C. magdalenae* | parcelas Humboldt |  | - |
| T2P4 | *C. magdalenae* | parcelas Humboldt |  | - |

**Table 2.** *Calliandra* species from Colombia that were included in this study.

|  |  | Souza et al. |  |
| --- | --- | --- | --- |
| Species |  | 2013 | This study |
| *Calliandra angustifolia* | Native | X |  |
| *Calliandra antioquiae* | Endemic |  | X |
| *Calliandra asplenoides* | Native | X |  |
| *Calliandra coriacea* | Native | X | X |
| *Calliandra falcata* | Introduced |  | X |
| *Calliandra fasciculata* | Native | X |  |
| *Calliandra glomerulata* | Native | X |  |
| *Calliandra guildingii* | Native | X |  |
| *Calliandra haematocephala* | Introduced | X | X |
| *Calliandra laxa* | Native | X |  |
| *Calliandra magdalenae* | Native | X | X |
| *Calliandra medellinensis* | Endemic |  | X |
| *Calliandra pittieri* | Native | X |  |
| *Calliandra purdiei* | Native | X | X |
| *Calliandra purpurea* | Native | X |  |
| *Calliandra riparia* | Introduced | X | X |
| *Calliandra surinamensis* | Native | X |  |
| *Calliandra tergemina* | Native | X | X |
| *Calliandra tolimensis* | Native |  | X |
| *Calliandra trinervia* | Native | X | X |
| *Calliandra tsugoides* | Native | X |  |

### DNA extraction, amplification, and sequencing

Total DNA was extracted from leaf tissues following the CTAB protocol adapted from Ivanova et al. (2006). The Internal Transcribed Spacer (ITS) encompassing *ITS1* and *ITS2*, was amplified using the primer pair *ITS*-5P (5’-GGAAGGAGAAGTCGTAACAAGG-3’) and *ITS*-4 (5’-TCCTCCGCTTATTGATATGC-3’) as designed by Möller and Cronck (2001) and White et al. (1990), respectively. The *trnL* intron and *trnL*-*trnF* intergenic spacer were amplified as a single PCR product using the primers *trnL*-c (5’-CGAAATCGGTAGACGCTACG-3’) and *trnL*-*trnF* (5’-ATTTGAACTGGTGACACGAG-3’), following the approach of Taberlet et al. (1991).

DNA amplification was performed using a master reaction mix with a final volume of 15 μL. This mixture contained2 μL of template DNA (∼10-50 ng), 1X Taq buffer ((NH_4_)_2_SO_4_), 200 μM of each deoxynucleoside triphosphate, 2 mM MgCl_2_, 0.2 μM of each primer, 0.4 μg/μL bovine serum albumin, and 1 U of Taq DNA polymerase (OneTaq, New England Biolabs). The PCR conditions included an initial denaturation step at 94°C for 3 min, followed by 35 cycles of denaturation at 94°C for 30 s, primer annealing (55°C for *ITS* and 50°C for trnL) for 40 s, and extension at 72°C for 60 s. Afinal extension step was performed at 72°C for 5 min. PCR products were visualized by electrophoresis on a 1.5% (w/v) agarose gel. Subsequently, both strands of the PCR products were sequenced using Sanger sequencing at the Universidad de los Andes in Bogotá, Colombia.

### Data edition, alignment, and phylogenetic analysis

We compiled a comprehensive molecular data matrix comprising 120 taxa, encompassing 117 as outlined by Souza et al. (2013) and the 3 new species sequenced here: this incorporated 142 *ITS* sequences and 146 *trnL* and *trnL-F* sequences, including different accessions of the same species. Assembly and refinement of complementary strands were conducted using Geneious v.6 (Kearse et al. 2012) with meticulous visual validation to ensure strand complementarity. Alignments were executed with MUSCLE v. 3.8425 (Edgar 2004). We treated gaps as missing data, and ambiguous segments were excluded from the alignments.

For each genetic marker, we evaluated the model of molecular evolution through ModelTest v.0.1.7 (Darriba et al. 2020). A maximum likelihood (ML) phylogenetic reconstruction was executed using 100 bootstrap replicates within the RAxML-HPC Blackbox program (Stamatakis 2014), implemented on CIPRES (2019) (Miller 2010). Bayesian phylogenetic analyses were run on MrBayes v.3.2 (Ronquist and Huelsenbeck, 2003). We conducted two independent runs, each spanning 30,000,000 generations and encompassing four chains (three heated and one cold). We employed a chain temperature of 0.2 and uniform priors. Trees were sampled every 1,000 generations, and a burn-in phase discarded the initial 25% of the runs. We used a majority-rule consensus to generate a Bayesian inference tree, featuring posterior probabilities (PP), from the two independent runs. The resulting trees were refined using Figtree v.1.4.4 (Rambaut 2016). To infer a phylogenetic network, we applied a neighbor-net algorithm using SplitsTree v. 5 (Huson & Bryant, 2005).

## RESULTS

We generated 80 novel sequences (34 for *ITS* and 46 for *trnL*-F) of *Calliandra* species occurring in Colombia, including multiple individuals from three *Calliandra* species absent in previous phylogenetic analyses: *C. medellinensis*, *C. tolimensis*, and *C. antioquiae* (Fig. 1, Table S1). *ITS* alignment had an average sequence length of 682bp. Among these, 61 sites provided informative insights under the parsimony principle, whereas 621 sites were conserved. Likewise, the final *trnL* and *trnL-F* alignment consisted of 146 sequences, with an average sequence length of 918. Of these, 92 sites were informative under parsimony and 826 sites were conserved.

The combined data matrix integrated 62 sequences, with an average sequence length of 1,090. Among these, 44 sites were informative, whereas 1,046 sites were determined to be non-informative based on parsimony.

### Nuclear region ITS

The optimal molecular evolution model identified for the *ITS* region was *TIM3+I+G4*. We achieved resolution of relationships among most species within the monophyletic *Calliandra* sect. *Androcallis*. Nevertheless, the tree did not fully recover the shallow phylogenetic relationships in this section (Fig. 3).

**Figure 3.**
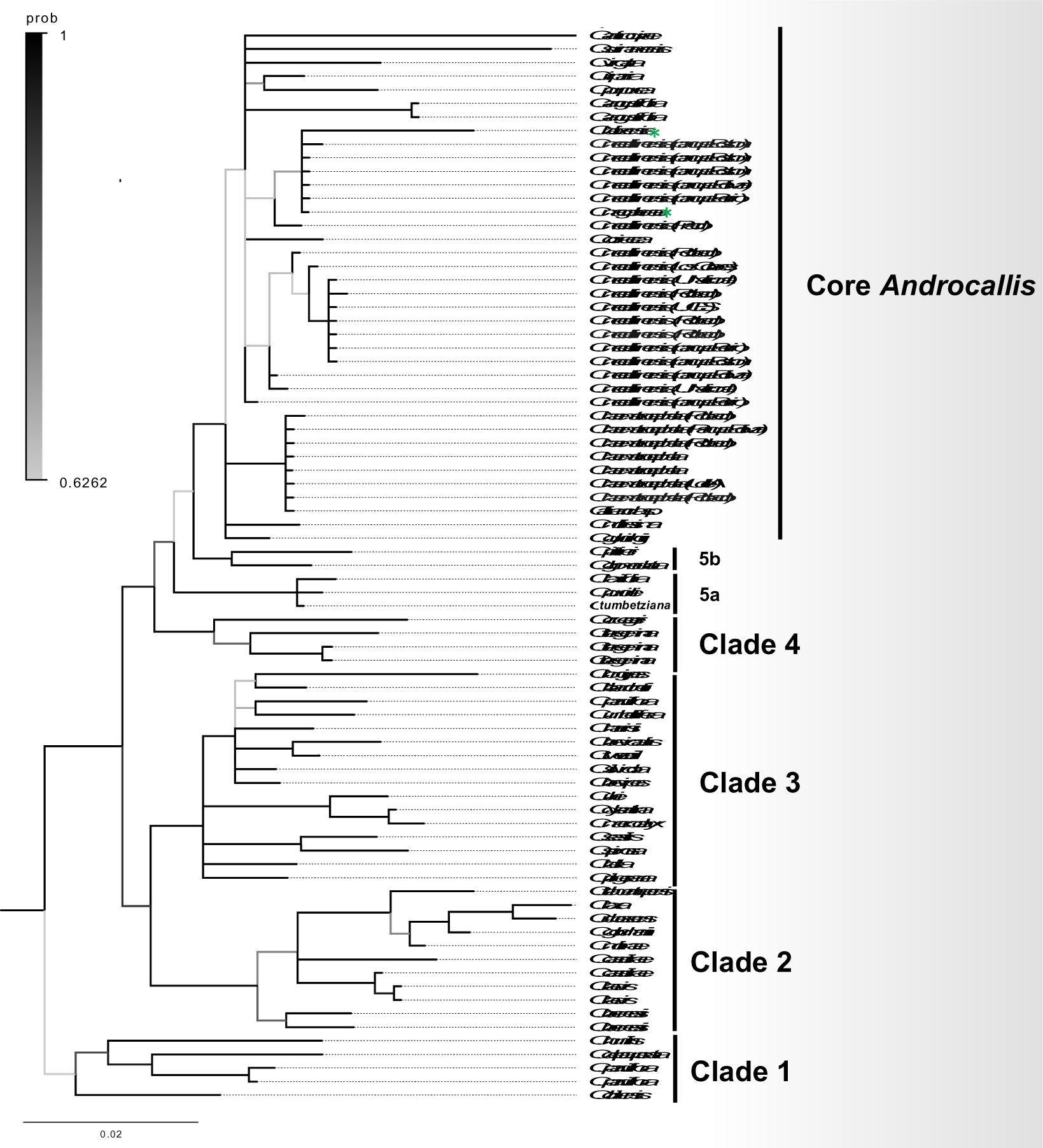
Bayesian posterior probability phylogeny of nuclear DNA (*ITS*) data for *Calliandra* sect*. Androcallis*. Species from *C.* sect. *Monticola* and *Microcallis* were used as outgroup. Black to grey bar corresponds to Bayesian inference posterior probabilities (PP) values. Bayesian inference (BI) consensus tree generated by MrBayes exhibited the highest score of –ln 4768.3362

The *Androcallis* phylogeny recovered six to seven distinct primary clades contingent upon the inclusion or omission of *C*. subsect. *Microcallis* within *C.* sect. *Androcallis*. Here, we recognized *C.* subsect. *Microcallis* as sister to *C.* sect. *Androcallis*. One early divergent clade, henceforth referred to as Clade 1, emerged as sister to the remainder of *Androcallis*. This clade includes species *C. humilis*, *C. parviflora*, *C. depauperata*, and *C. chilensis* (PP: 0.63). The remaining *Androcallis* species comprise four well-supported clades.

Clades 2 and 3 (PP:0.90) were found to be sister to clade 4 and 5 (PP:0.99). Clade 2 includes species *C. brenesii*, *C. laevis*, *C. caeciliae*, and a robust subclade including species *C. molinae*, *C. goldmanii*, *C. rubescens*, *C. laxa*, and *C. tehuatepecensis* (PP: 1). Relationships among species in Clade 3 are poorly resolved and comprise species *C. pilgerana* and *C. bella,* forming a polytomy with three other subclades: *C. sessilis* + *C. spinosa* (PP: 1), *C. ulei* (*C. macrocalyx* + *C. dysantha*) (PP: 1), and *C. brevipes*, *C. silvicola*, *C. teewedii*, *C. brevicaulis*, *C. harrisii*, *C. umbellifera*, *C. parvifolia*, *C. blanchetii*, *C. longipes* (PP: 1). Species from Clade 4 comprise *C. tergemina* and *C. cruegeri* (PP: 0.97), while Clade 5 contains three subclades 5a (5b + 5c + core *Androcallis*) (PP: 1): Subclade 5a includes *C. tumbetziana*, *C. purdiei*, and *C. taxifolia* (PP: 1). Subclade 5b comprise*s C. pittieri* and *C. glomerulata* (PP =0.99). The last subclade, 5c, is denoted here as core *C.* sect. *Androcallis* (PP: 0.96; Fig. 3; Table 1).

Among core *Androcallis,* backbone relationships are not well supported except for a clade formed with several accessions of *C. haematocephala*, and a second one including some of the *C. medellinenses* accessions from parks and neighborhoods around Medellín (Fig.1, Fig. 3). Although the evidence is not conclusive, it suggests that *C. medellinensis* is not monophyletic, and some accessions growing in parks around the center of Medellin are more closely related to *C. magdalenae* and/or *C. haematocephala* (PP:0.7) (Fig. 3 and 4).

**Figure 4.**
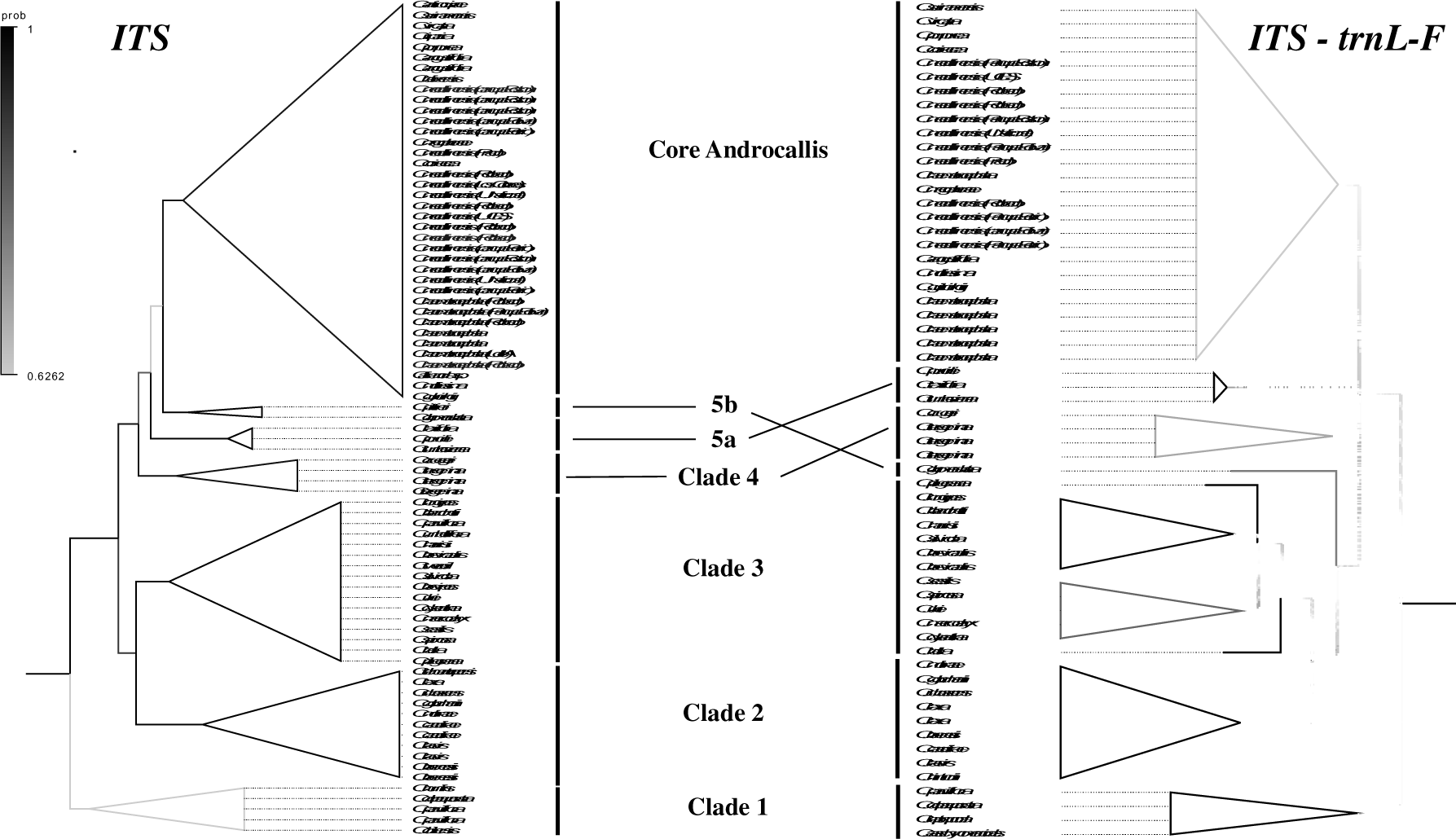
Comparison of Bayesian phylogenetic topologies of two trees: Combined nuclearDNA (*ITS*) and cpDNA (*trnL* - *trnL-F*) and nuclear DNA (*ITS*) data for *Calliandra* sect. *Androcallis*. Species from sect. Monticola and *Microcallis* were employed as outgroup. The gradation from black to grey bars corresponds to the Bayesian inference posterior probabilities (PP) values.

### Chloroplast region trnL and trnL-F

The best molecular evolution model for the *trnL* and *trnL-F* region was TIM1+I+G4. Support values from Maximum Likelihood Bootstrap (MLB) were sometimes markedly lower than BI posterior probability values for the same clade, generally supported by relatively few characteristics. The ML and BI analyses based on one cpDNA region yielded nearly non-conflicting topologies, with BI resolving and better supporting clades that were in a polytomy or poorly supported in ML analysis. A summary of cpDNA phylogenetic results is shown in Suppl. Fig. 1. *Androcallis* was recovered as monophyletic and all species from Colombia sequenced here were included within this clade. The backbone of this phylogeny was not resolved by ML and BI analyses; however, small clades were recovered.

Phylogenetic relationships using the *trnL* and *trnL-F* region were poorly resolved here. Clade 1 and 5a from the *ITS* phylogeny (Fig. 3, Suppl. Fig. 1) are strongly supported here even though not all same species are included in both phylogenies. A clade containing several accessions of *C. medellinensis* (Suppl. Fig. 1), *C. haematocephala*, and *C. magdalenae* was moderately supported here. Only one accession of *C. medellinensis* from Mariquita seems to be more closely related to a *C. tergemina* accession collected in a neighborhood south of Medellín (Terminal del Sur). This phylogenetic proposal suggests that *C. antioquiae* is monophyletic, whereas *C. magdalenae* and *C. haematocephala* are non-monophyletic, as accessions of both species are in different clades (Suppl. Fig. 1).

### Analysis of the combined data set

The optimal molecular evolution model for the combined dataset was GTR. In this phylogeny, only clades 1, 2, 3, and 5a, as identified in our *ITS* phylogenetic tree, were recovered with moderate-to-robust support (Fig. 4). However, the overall resolution of the phylogenetic backbone remained partial. Clade 1 encompasses *C. parviflora*, *C. depauperata*, *C. leptopoda*, and *C. aeschynomenoides*, strongly supported as the sister group to the remainder of *C.* sect. Androcallis. Clade 2 emerged as sister to Clade 3. However, the comprehensive arrangement of sect. The *Androcallis* backbone provided clear resolution using combined nuclear and chloroplast markers (Fig. 4).

Clade 2 comprises *C. molinae*, *C. goldmanii*, *C. rubescens*, *C. laxa*, *C. brenesii*, *C. caeciliae*, *C. laevis*, and *C. hintonii*. The phylogenetic relationships among these species within this clade were fully resolved. Clade 3 includes *C. sessilis*, *C. spinosa*, *C. ulei*, *C. macrocalix*, *C. dysantha*, *C. longipes*, *C. blanchetti*, *C. harrisii*, *C. silvicola*, *C. brevicaulis*, *C. pilgerana*, and *C. bella*. The analysis also uncovered three additional clades: Core Androcallis, Clade 5a containing *C. purdiei*, *C. taxifolia*, and *C. tumbeziana*, and Clade 5b represented by only *C. glomerulata*. Notably, the relationships between clades 2+3, 4, 5a, 5b, and core Androcallis in this phylogeny lacked robust support (Fig. 4).

### Core *Androcallis* phylogenies

To achieve a finer resolution of the relationships among species within the Core *Androcallis* group, we conducted targeted subsampling of species from the broader *Androcallis* phylogeny and employed Clades 5a and 5b as outgroups. This focused analysis substantially enhanced the resolution of the core *Androcallis* phylogeny, except for certain species exhibiting limited molecular variation within both *the ITS* nuclear region and *TrnL*-*F* plastid region (Figs. 4 and 5). Support values obtained through MLB analysis were noticeably lower than the BI posterior probability values for the same clades, primarily because a relatively limited number of informative characters (Suppl. Fig. 2-3). BI analysis, in contrast, succeeded in resolving and providing better support for clades that had previously appeared as polytomies or received weak support in the ML analysis.

**Figure 5.**
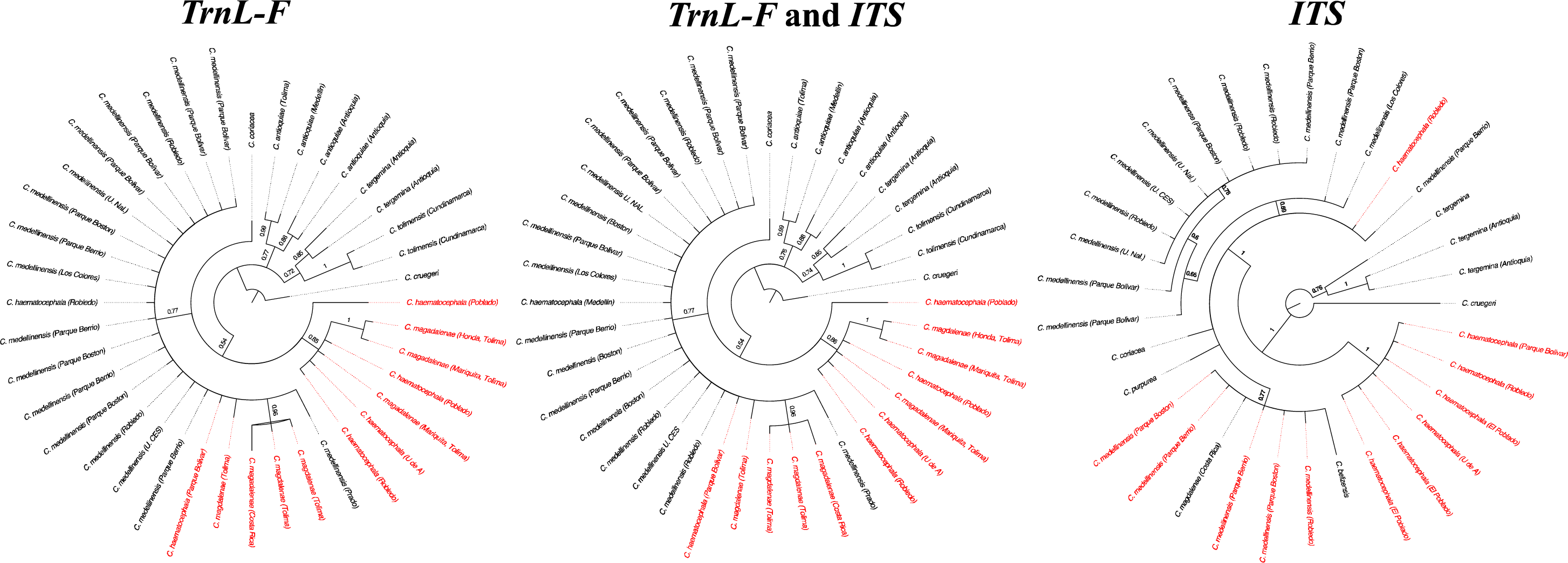
Comparison of Bayesian phylogenetic topologies, a plastid phylogeny cpDNA (*trnL-F*), a combined nuclear DNA (*ITS*) and cpDNA (*trnL* and *trnL-F*) phylogeny, and nuclear DNA (*ITS*) phylogeny for Core *Androcallis*, with emphasis in *Calliandra medellinensis*. For the outgroup, species from clade 4 were employed. Bayesian inference posterior probabilities (PP) values are indicated in branches. Besides *C. medellinensis*, taxa flagged in red appeared to be non-monophyletic.

#### Nuclear region ITS

Within the Bayesian phylogenies, PP substantiates the hypothesis of non-monophyly for both *C. haematocephala* and *C. medellinensis* (Fig. 5). Accessions of *C. haematocephala* sampled from various public parks around Medellin formed a well-supported clade positioned as the sister group to the remainder of the species within the core *Androcallis* (PP:1). This sister clade, in turn, encompasses a polytomy consisting of *C. purpurea*, *C. coriacea*, *C. medellinensis*, and an accession of *C. haematocephala*. It also includes a moderately supported clade containing *C. belizensis* and *C. magdalenae* from Costa Rica, which were both sequenced by de Souza et al. in 2013. This clade also comprises several accessions of *C. medellinensis* (PP:0.77) (Fig. 5). When examining the alignment of *ITS* sequences, two SNPs observed at positions 354 and 554 represented a transitional mutation corresponding to a guanine (G) shared by all species and accessions mentioned above (Fig. 5, Suppl. Fig. 2). In addition, a single ambiguous nucleotide (R: A/G) at position 354 distinguishes one accession of *C. haematocephala* from Robledo from other accessions of the same species, including one from the same public park in Medellín. Interestingly, this ambiguous nucleotide at position 354 was shared by two *C. medellinensis* accessions collected from Parque de Boston and Parque Berrío (Suppl. Fig. 2). Within the remaining *C. medellinensis* group in this polytomy, a second moderately supported clade showed partial resolution and was characterized by several accessions of *C. medellinensis* (PP:0.69), all sharing an adenine (A) at the same position (Fig. 5, Suppl. Fig. 2).

#### Chloroplast region trnL and trnL-F

Within the Bayesian phylogenetic analysis, the PP support lends strong evidence to the hypothesis of non-monophyly for three species: *C. magdalenae*, *C. haematocephala*, and *C. medellinensis* (Fig. 5). At the root of this phylogenetic tree, a polytomy was observed, encompassing an accession of *C. haematocephala* from El Poblado Medellín, *C. coriaceae,* and four moderately supported clades. The first clade was comprised of multiple accessions of both *C. magdalenae* and *C. haematocephala* (PP:0.85). The second clade includes all *C. medellinensis* accessions without any resolution, along with an accession of *C. magdalenae* from Tolima and *C. haematocephala* from Parque Bolívar in Medellín, as well as a clade containing several *C. magdalenae* accessions from Tolima and Costa Rica. The latter was sequenced by Souza et al. (2013). The remaining two clades within the backbone polytomy were completely resolved: One formed by *C. antioquiae* accessions (PP:0.77), and the other by *C. tolimensis* accessions (PP:0.72) (Fig. 5).

Examination of the alignment revealed that all *C. magdalenae* accessions shared an insertion/deletion at nucleotide 1046, except for one accession from Tolima, which fell outside the clade formed by the other *C. magdalenae* accessions from Tolima and Costa Rica. Additionally, a SNP (C-T) at position 1066 in the alignment is shared among all *C. haematocephala* accessions, some *C. magdalenae* accessions, as well as *C. antiquiae* and *C. tolimensis* accessions (they share a C at this position). Conversely, the clade containing all *C. medellinensis* accessions and other *C. magdalenae* accessions, along with one accession *of C. haematocephala*, shared a T at this position (Suppl.Fig. 3).

#### Analysis of the combined dataset

This phylogeny showed the same topology with similar posterior probabilities and bootstrap values when compared to the plastid phylogeny (Fig. 5).

## DISCUSSION

In this study, we elucidated the backbone relationships within *Calliandra* sect. *Androcallis*, thereby establishing a phylogenetic framework that can serve as a valuable reference for forthcoming investigations pertaining to taxonomy, biogeography, and character evolution within this group. This phylogeny further advances previous results published by de Souza et al. (2013). Our approach also underscores the significance of incorporating both chloroplast and nuclear data while ensuring a comprehensive geographic representation of specimens to resolve phylogenetic intricacies in *Calliandra*. Within *Calliandra* sect. *Androcallis*, our analyses revealed the presence of at least five primary clades. Nevertheless, several pivotal questions regarding the evolutionary relationships among the core *Androcallis* species, particularly those collected in Colombia, remain unanswered (Figs. 3-5).

### A resolved *Androcallis* backbone

The phylogenetic hypothesis based on nuclear *ITS* and chloroplast *trnL-F* sequences corroborated previous proposals, which demonstrated the monophyly of the genus *Calliandra* sect. *Androcallis* + *Microcallis* (Souza et al. 2013). The monophyly of this clade is also confirmed by their pollen morphology; species from this group lack an apical appendage in pollen grains found in *Calliandra* section *Monticola* (Santos and Romao 2008, de Paula et al. 2020). While only the polyad size distinguished *Calliandra* sect. *Androcallis* from *Microcallis*, the smallest polyad size are present in the *Microcallis* (de Paula et al. 2020). The phylogeny of combined nuclear and plastid markers also recovered the five clades identified using *ITS*, further reinforcing the robustness of our results (Fig. 4).

Our expanded dataset introduced samples from Colombian species *C. medellinensis*, *C. antioquiae*, and *C. tolimensis* for the first time. Additionally, we incorporated new sequences for various species collected in Colombia, including *C. haematocephala*, *C. riparia*, *C. magdalenae*, *C. coriaceae*, *C. trinervia*, *C. tergemina*, and *C. purdiei*. While the *ITS* molecular marker proved instrumental in elucidating the *Androcallis* backbone, it did not provide comprehensive resolution of relationships among core *Androcallis* species, including those from Colombia. The *Androcallis* group comprises widely distributed species extending from Argentina to southern United States. They exhibit diverse growth habits, ranging from bushy to small trees or subshrubs, and often exhibit rhizomatous tendencies. They also feature lateral inflorescences in brachyblasts or terminal umbels (Barneby 1998).

Barneby’s comprehensive classification of *Calliandra* in 1998 identified five sections, with a particular emphasis on inflorescence architecture as a defining trait. However, our study focuses exclusively on species from the *Microcallis* and *Androcallis* sections. Barneby differentiated *Microcallis* species primarily by the presence of a terminal pseudoraceme of heads or umbels, a feature also found in *C.* sect. *Calliandra*, a group not included in this study. Additionally, he further separated *Microcallis* species based on perianth dimensions. Notably, in both, our study, and Souza et al. (2013), *Microcallis sensu* Barneby did not form a monophyletic group. Instead, it included a mixture of species from Barneby’s *Microcallis* and some from the *Androcallis* section. This raises questions about the utility of inflorescence architecture as a reliable systematic character in this context. Our findings suggest that species from *Microcallis* are sister to those from *Androcallis*, implying a basal position for *Microcallis* within *Androcallis* (Clade 1) (Fig. 3-5).

The series proposed by Barneby (1998) within his taxonomic classification of *Androcallis* does not align with phylogenetic reconstructions, indicating that morphological characters at the series level do not provide informative insights. However, geographical patterns emerge as a valuable indicator of evolutionary history. For instance, species belonging to Clade 2 receive strong support, corroborating findings by Souza et al. (2013). Nevertheless, the relationship of this clade with the remainder of *Androcallis* species is reinforced solely by our study, where it is identified as the sister clade to the rest of *Androcallis* species (Figs. 3 - 4). Clade 2 predominantly comprises species from Central America, except for *C. laxa*, which extends its distribution into South America.

Clade 3, encompassing species *C. spinosa* and *C. sessilis*, is robustly supported as sister group to Clade 2 in our *ITS* phylogeny (Fig. 3). The phylogenetic placement of *C. spinosa* and *C. sessilis* remained unclear in de Souza et al. (2013), whereas in our phylogeny both species are included in Clade 3. Notably, species within Clade 3 predominantly inhabit southern South America, including regions of Peru, Bolivia, Chile, Paraguay, and several are confined to specific geographic areas in Brazil. Polyploidy has been detected in several species of *Androcallis*, particularly in species of Clade 3 (de Paula et al. 2020).

Clade 4 is strongly supported as the sister to the remaining *Androcallis* species (Clade 5) and exhibits a distribution range spanning from Venezuela to northern Brazil and Peru within the Amazonas Forest region. Additionally, Clade 5 has been further subdivided into smaller subclades, including *C. taxifolia*, *C. tumbetziana*, and *C. purdei* (5a), as well as Clade 5b, comprising species *C. pittieri* and *C. glomerulata*, distributed in Northern South America, including Ecuador, Colombia, Venezuela, northern Brazil, and Peru. Most species with distribution in Colombia included in this study are placed in *Androcallis* Clade 5, the majority in core *Androcallis* (Fig. 3) (Bello and Forero 2005). In general, the overall phylogenetic resolution within the core *Androcallis* group was suboptimal with the nuclear and chloroplast markers employed in this study.

These phylogenetic insights underscore the significance of geographic distribution as critical factors in understanding the evolutionary relationships and diversification within *Androcallis*. While the *ITS* phylogeny exhibited better resolution, discerning strongly supported subclades, certain inconsistencies arose, particularly in the data sets where only one marker was available per species (Fig. 5). For instance, in the combined phylogeny, Clade 1, also known as *Microcallis,* garnered substantial support as it emerged as the sister group to the remaining *Androcallis* species, a distinction less pronounced in the *ITS-*based topology. Conversely, Clade 4 received robust support in the *ITS*-based topology as the sister group to Clade 5, whereas the combined markers topology depicted a different relationship (Fig. 4, 5).

### Core Androcallis and Calliandra medellinensis

Notably, our analysis indicated that *C. haematocephala*, *C. magdalenae*, and *C. medellinensis* did not exhibit monophyly across all phylogenetic assessments. These three taxa also represented challenges for their taxonomic identification since they are very similar to one another. Bello and Forero (2005) distinguished several varieties in *C. magdalenae* based on morphological variation of folioles and number of pinnes. It is important to highly these so-called species are mostly distinguished from one another morphologically by the number of pinates and folioles present in leaves, type of indument in branches, variation in the number of stamens and flower color (Bello and Forero 2005).

The species *Calliandra medellinensis* is an endemic taxon from Colombia that is only known from planted individuals in Medellín, Antioquia, and a few rare collections from Tolima and Cundinamarca. This taxon has been a subject of taxonomic interest, with some taxonomists (Bello and Forero 2025) previously suggesting its hybrid nature due to morphological similarities with *C. haematocephala*. Our findings suggest a complex evolutionary history involving reticulation between *C. haematocephala* and *C. magdalenae*. However, this should be pondered with caution because the absence of clear phylogenetic signals in the selected markers and the selective marker amplification of Colombian samples. This needs further confirmation, with a complete taxon sample for *Androcallis*, more informative molecular markers, from both maternal and biparental inheritance, and karyological studies for these species.

Reticulate evolutionary scenarios, encompassing processes such as chloroplast capture and hybridization, may explain the observed discrepancies between chloroplast and nuclear DNA data (Tsitrone et al. 2003; Linder and Rieseberg 2004). It is imperative, however, to exercise caution when attributing these discrepancies solely to specific causes, as other stochastic factors may yield analogous patterns. Incomplete lineage sorting, for instance, is another plausible explanation, although it is typically invoked for groups displaying intricate organellar DNA relationships within and among species that are not generally sympatric.

The species *C. medellinensis* samples were obtained from the metropolitan area of Medellín, comprising a population of 26 individuals that exhibited varying degrees of phenotypic variation in vegetative and floral characteristics. Reports indicate that *C. medellinensis* has been cultivated as an ornamental species in Medellín’s parks and streets. An unpublished study on the reproductive biology of *C. medellinensis* and *C. haematocephala*, conducted in Medellín, revealed significant differences in fecundity between the two species (Cano et al, 2017, *unpublished results*). While *C. haematocephala* exhibited higher nectar rewards, more floral visitors, increased pollen loads, and greater fruit setting, *C. medellinensis* did not. This reduced fruit and seed setting in *C. medellinensis* raises the possibility that it represents a poorly adapted and recent hybrid. It is equally plausible that *C. medellinensis* could be self-incompatible and cross-pollination rate among accessions be low. This could complicate conservation efforts, particularly in urbanized environments where human activities may disrupt interspecific interactions contributing to pollination and dispersion. However, if indeed a hybrid, *C. medellinensis* may possess floral visitors and effective pollinators that are generalists, visiting other species, including parental ones.

Furthermore, the habitat fragmentation among *C. medellinensis* populations hinders genetic exchange and population expansion, exacerbated by the apparent infertility of the seeds and the reliance on vegetative propagation methods such as cuttings and laboratory-based tissue culture. Despite these challenges, assessing the extinction risk of *C. medellinensis* under IUCN Red List protocols remains inconclusive since IUCN does not typically evaluate hybrid taxa, except for apomictic plant hybrids.

In conclusion, this study contributes to the phylogenetic elucidation of the *Calliandra* sect. *Androcallis* phylogeny providing an evolutionary framework in which to test ecological, biogeographical among other hypotheses. Notably, *C. medellinensis*, *C. antioquiae*, and *C. tolimensis* were included in *Calliandra* sect. *Androcallis*, particularly within Clade 5 subclade Core *Androcallis*. However, the non-monophyly observed in *C. medellinensis* necessitates further validation through informative molecular markers to elucidate the potential occurrence of interspecific hybridization between *C. haematocephala* and *C. magdalenae*, its timing, and the taxa involved.

## ACKNOWLEDGMENTS

We thank Mónica Marcela Gaviria, Juan Sebastián Pino, and Laura Victoria Cano for their contribution to the initial formulation of this project and Carolina Castellanos Castro and Luis Eduardo Mejía for their support to conduct field work. We also thank members of the Comparative Biology Laboratory of the Center for Biological Research (CIB Medellín), the Alexander von Humboldt Institute, and the Animal Genetics Laboratory of the University of Antioquia for their assistance in the lab and analyzing data as well as to researchers and staff of the Joaquín Antonio Uribe Herbarium (JAUM), the University of Antioquia Herbarium (HUA), and the National University Herbarium at Medellín (MEDEL).

## Author contributions statement

TAG: Co-developed questions and framework, performed analyses, mentored student author, assisted with obtaining funding, wrote and edited text.

JDS: Co-developed questions and framework, performed analyses, wrote text.

HA: Co-developed questions, obtained funding, made collections, edited text.

IDSC: Co-developed questions and framework, mentored student author, assisted with obtaining funding, edited text.

With contributions of all authors, to the analysis and text.

## Declarations

We have all collection permits required by the Colombian government to collect leaves samples of mature individuals for DNA extractions: Autoridad Nacional de Licencias Ambientales (ANLA)-8 de Octubre 2015, Resolución Número 1263.

All methods were carried out in accordance with relevant guidelines and regulations.

## Availability of data and materials

All data have been deposited in Bioproject (XXXX)

## Consent to participate

Not applicable

## Consent for publication (include appropriate statements)

Not applicable

